# Pan-cancer transcriptomic analysis reveals a tissue-specific loss of identity

**DOI:** 10.1101/2022.12.12.520045

**Authors:** Gabriel Arantes dos Santos, Kasit Chatsirisupachai, Roberto A. Avelar, João Pedro de Magalhães

## Abstract

Understanding changes in cell identity in cancer and ageing is of great importance. In this work, we analyzed how gene expression changes in human tissues are associated with tissue specificity during cancer and ageing using transcriptome data from TCGA and GTEx. We found a loss of cell identity during ageing in around 40% of tissues. In cancer, we observed a pattern of downregulation of genes specific to the tissue of origin of the tumor for most cancer types. Moreover, we observed an activation of genes not usually expressed in a tissue as well as an upregulation of genes specific to other tissues in cancer. These patterns in cancer were associated with patient survival. The age of the patient, however, does not influence these patterns. Thus, we identified a moderate but statistically significant pattern of loss of cell identity in ageing, and a clear pattern in cancer, where during tumorigenesis, cells express genes specific to other organs while suppressing the expression of genes from their original tissue. The loss of cellular identity observed in cancer is associated with prognosis and is not influenced by age, suggesting that it is a crucial stage in carcinogenesis.

## Introduction

Tissues are formed by the combinations of cells expressing different transcripts and proteins, which shape their morphology and function (1). Differences between tissues are driven by transcriptomic programs and signatures, which change during normal organismal development and also in detrimental processes such as disease and ageing (2, 3).

One hypothesis in gerontology is that tissues lose their cellular identity during ageing, which contributes to age-related dysfunctions. Although recent studies support this idea for some tissues, we still do not have enough evidence to confirm this hypothesis, which highlights an important topic for the field (4-6).

Cancer cells are known to gain plasticity and stemness during tumor initiation and progression, and recently, unlocking phenotypic plasticity has been considered a “new” cancer hallmark (7, 8). Ageing is one of the main risk factors for most cancers, which may link tissue specificity processes, the onset of cancer and progressive age-related disruptions (9). Moreover, cancers of different origins have different behaviour and development, and the relationship between the tumor and the original tissue needs to be better elucidated to improve our understanding of cancer biology and find better treatment options (10-12). Finally, growing evidence confirms that the molecular landscape of cancers from old and young patients are different, but we do not yet know whether patterns of tissue-specific identity influence these differences in any way (13).

In this study, we explore the relationship between genes differentially expressed in cancer and ageing with tissue-specific identity, using data from TCGA (The Cancer Genome Atlas), a comprehensive consortium that uses thousands of cancer samples to try to decipher the tumour landscape through sequencing and clinical data, and GTEx (Genotype-Tissue Expression), a public resource to study tissue-specific gene expression and regulation from healthy samples.

## Methods

### Data Acquisition

The mRNA expression data in read counts from the TCGA harmonized data (data aligned to hg38) and clinical data (XML files) were downloaded using TCGAbiolinks (version 2.14.1), as described in Chatsirisupachai et al (14, 15).

RNA-Seq-based gene expression data from normal tissues (version 8) were downloaded from the GTEx portal (https://gtexportal.org) (16). According to the GTEx documents, the raw sequence reads were aligned to the human reference genome GRCH38/hg38.

First, we separated the TCGA or GTEx data according to cancer/tissue, and then we removed genes with less than 1 count in more than 30% of samples. TCGA and GTEx analyses were done independently. We used the biomaRt package to keep only the protein-coding genes in all analyses (17).

### Genes differentially expressed in ageing (ageing-DEGs)

GTEx data was used to find differentially expressed genes with age. Samples without complete information were filtered out, and tissues with less than 50 complete samples were also excluded. GTEx only lists age ranges (i.e., 20-29, 30-39, 40-49, 50-59, 60-69 and 70), so age was approximated to 25, 35, 45, 55, 65 and 75, respectively (3). The sample numbers for each tissue used in this study are shown in table S1.

GTEx GENCODE gene IDs were converted to Ensembl gene IDs using the cleanid() function from the grex package version 1.9 (18). After ID conversion, 18 Ensembl IDs were duplicated – these genes were removed from the read counts.

Samples were grouped into their respective tissues and processed together. Read counts were first converted to counts per million (CPM) using the cpm() function with default parameters from the edgeR package(19-21). To correct for library size variation between samples, the trimmed mean of M-values (TMM) normalisation method was applied using the calcNormFactors() function with default parameters (22). Counts were then voom transformed to adjust for heteroscedasticity using the voom() function with default parameters (23)

For each tissue, FC with age was calculated using the model below. If any variable is not present in the tissue (e.g., sex for prostate or region for blood), it is disregarded in the analysis

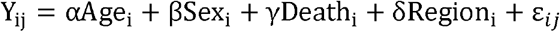

The variables are defined as follows:

- *Y*_*ij*_: The expression level of gene *j* in sample *i*.
- *Age*_*i*_: The age of sample *i* – continuous variable.
- *Sex*_*i*_: The sex of sample *i –* categorical variable.
- *Death*_*i*_: The death classification of sample *i* based on the 4-point Hardy scale *–* categorical variable (24).
- *Region*_*i*_: The tissue region cells were extracted from for sample *i –* categorical variable.
- ε_*ij*_: The error term for gene *i* in sample *j*.

Linear models were generated using the R package limma, using the lmFit() function with default parameters(25, 26). Genes were considered significant DEGs if they matched the following criteria: i) The p-values derived from the empirical Bayes moderated t-statistics were less than 0.05 after Benjamini-Hochberg (BH) FDR correction; and ii) the absolute log2 (fold change) across 50 years of age (from 25 – 75), represented as 50* log2 (fold change) was greater than log2(1.5) (3).

### Genes differentially expressed in cancer (cancer-DEGs)

Of all the cancers available in TCGA, we selected for our analyses only projects with at least ten samples of adjacent normal tissue available. The selected TCGA projects are: BLCA, BRCA, COAD, ESCA, HNSC, KICH, KIRC, KIRP, LIHC, LUAD, LUSC, PRAD, READ, STAD, THCA and UCEC. For BRCA, we kept only samples from female patients. The sample numbers for each project (cancer and normal samples) are shown in table S1.

To generate a list of differentially expressed genes in cancer (cancer-DEGs), we compared tumour samples with adjacent normal tissue samples. Data were processed and analyzed using the R package DESeq2 (27) using default parameters. A gene was considered differentially expressed when Fold Change >2 and FDR <0.01. The p-value adjustment was made using BH methodology.

### Classification of genes based on specificity category and overlap analysis

First, we need to classify all genes according to their tissue specificity. For this, we downloaded the Tau Index-based classification by Daniel Palmer et al (28). Briefly, the tau index is based on GTEx data and indicates how specific or widely expressed a gene is, with tau = 1 indicating expression specific to only one tissue and tau = 0 indicating similar expression in all tissues.

We then created four specificity categories that were separated into two groups. The first group we called “Pan-tissue categories” is based only on tau values; therefore, these genes are the same for all the tissues analyzed. The second group we called “Tissue-Specific categories,” is based on tau’s values and tissue-specific average expression; therefore, the genes differ from tissue to tissue. The categories are shown below:

- Pan-tissue categories: “High Specificity genes” = Tau > 0.8; “Low Specificity genes” = Tau < 0.2
- Tissue-Specific categories: “Tissue-Specific genes” = Tau >0.95 and average expression >1 in the tissue of interest; “Tissue-Unexpressed genes” = Average expression = 0 in the tissue of interest

It is important to note that the Pan-tissue categories define how specific the expression of a gene is without specifying in which organ it is being expressed. On the other hand, Tissue-specific categories are based on the expression (or lack of expression) of genes only in the tissue analyzed.

We initially tried to analyze genes that are not expressed in any GTEx tissue, but observed minimal overlaps and only insignificant results. For this reason, we excluded these genes from further analysis. Thus, we kept only protein-coding genes and excluded transcripts where tau = NA (i.e., that are not expressed in any tissue). The values of the “Pan-tissue” group are based on the original paper by Palmer et al (28). In the “Tissue-specific” group, we tried to be as strict as possible to ensure tissue specificity, so we used extreme tau values. The “Tissue-Specific” category, we used tau> 0.95 because using tau= 1 would generate insufficient genes for further analysis, but we guaranteed tissue specificity and consistent expression using only genes with average expression above 1. The Tau data and average tissue expression used in this study is in table S2. The numbers of genes in each category and the background list (all remaining genes in the tau classification) are shown in Table 1.

**Table 1.** Number of DEGs and genes on specificity categories. *”Up” and “Down” represent whether genes are upregulated or downregulated

| High Specificity Genes | 4851 |  |  |  |  |  |
| --- | --- | --- | --- | --- | --- | --- |
| Low Specificity Genes | 3464 |  |  |  |  |  |
| Genes on background | 17836 |  |  |  |  |  |
|  | Up_Ageing | Down_Ageing | Up_Cancer | Down_Cancer | Tissue-unexpressed | Tissue-specific |
| Breast - BRCA | 163 | 42 | 1802 | 1379 | 3833 | 35 |
| Colon - COAD | 93 | 1256 | 1630 | 1710 | 3861 | 73 |
| Esophagus - ESCA | 102 | 90 | 1145 | 840 | 3958 | 37 |
| Kidney - KICH | 0 | 0 | 1848 | 1997 | 4238 | 113 |
| Kidney - KIRC | 0 | 0 | 2214 | 1269 | 4238 | 113 |
| Kidney - KIRP | 0 | 0 | 1809 | 1358 | 4238 | 113 |
| Liver - LIHC | 12 | 21 | 1476 | 882 | 5872 | 162 |
| Lung - LUAD | 279 | 230 | 1919 | 1518 | 3473 | 117 |
| Lung - LUSC | 279 | 230 | 2627 | 2176 | 3473 | 117 |
| Prostate - PRAD | 1182 | 1116 | 608 | 1097 | 3492 | 29 |
| Stomach - STAD | 25 | 16 | 1065 | 1591 | 4195 | 46 |
| Thyroid - THCA | 94 | 35 | 1167 | 641 | 3764 | 78 |
| Uterus - UCEC | 1024 | 1043 | 2364 | 1769 | 4122 | 27 |
| BLCA | - | - | 1415 | 1450 | 3928 | 44 |
| HNSC | - | - | 1290 | 1361 | - | - |
| READ | - | - | 1737 | 1748 | - | - |
| Adipose Tissue | 193 | 193 | - | - | 4223 | 27 |
| Adrenal Gland | 320 | 131 | - | - | 4736 | 56 |
| Blood | 280 | 143 | - | - | 5988 | 126 |
| Blood Vessel | 764 | 153 | - | - | 4481 | 25 |
| Brain | 430 | 360 | - | - | 3899 | 356 |
| Heart | 81 | 36 | - | - | 5483 | 51 |
| Muscle | 281 | 99 | - | - | 6289 | 83 |
| Nerve | 299 | 378 | - | - | 3934 | 77 |
| Ovary | 371 | 409 | - | - | 4303 | 42 |
| Pancreas | 58 | 20 | - | - | 5610 | 67 |
| Pituitary | 9 | 2 | - | - | 3382 | 242 |
| Salivary Gland | 951 | 526 | - | - | 3899 | 94 |
| Skin | 11 | 34 | - | - | 3874 | 195 |
| Small Intestine | 284 | 690 | - | - | 3688 | 119 |
| Spleen | 34 | 12 | - | - | 4277 | 119 |
| Testis | 102 | 36 | - | - | 1961 | 1248 |
| Vagina | 214 | 88 | - | - | 1588 | 46 |

After that, we overlapped the DEGs (cancer and ageing) with the four categories using basic R functions. The overlap was considered significant if FDR < 0.05 (Fisher’s exact test followed by Benjamini-Hochberg correction).

To make sure that the pattern found was biologically accurate, we used an alternative specificity classification and repeat the overlaps with the same parameters. For this, we downloaded data from Uhlén et al., where the authors’ classified genes based on RNA expression in a tissue-specific manner (29). Briefly, in this study, genes are divided into six main categories: “Tissue enriched”, “Tissue enhanced”, “Group enriched”, “Expressed in all”, “Mixed” and “Not Detected”. We then adapted these categories to our study, where we have the following “alternative categories” (number of genes and background list in each category in table S3):

- Pan-tissue categories: High specificity genes = “Tissue enriched”, “Tissue enhanced” and “Group enriched”; Low specificity genes = “Expressed in all” and “Mixed”,
- Tissue-Specific categories: Tissue-Specific genes = “Tissue enriched”, “Tissue enhanced” or “Group enriched” in the tissue of interest Tissue-unexpressed = FPKM < 1 in the tissue of interest

It is important to note that in Palmer’s classification, HNSC and READ data could not be analyzed in a tissue-specific manner as we do not have data available for the corresponding normal tissue (tonsil and rectum). In the alternative classification, BRCA, UCEC, blood, blood vessel, breast, nerve, pituitary, uterus, and vagina face the same limitation since their respective tissues are unavailable.

### Survival analyses

For this analysis, we selected only the genes that lie in the overlap between the cancer-DEG and one of the four categories. We use the expression signature of those genes to construct overall and disease-free survival curves. All survival analyses were performed on GEPIA2 using the median expression of the signature to segregate the two groups (30). The constructed heatmap was based on the log2 hazard ratio (Mantel-Cox test), and the result was considered significant when FDR <0.1.

### Cancer analyses based on patients’ age

To analyze whether the patient’s age influenced the loss of tissue identity in cancer, we separated the TCGA samples into two groups: young and old. The young group comprises 30% younger samples and the old group 30% older samples. We kept only cancers with at least ten control samples in each normal group (old and young normal). To avoid confounding factors, we compared the two groups regarding the T pathological stage (Fisher’s exact test); if the p-value <0.05, we considered the groups different and excluded this cancer from the subsequent analysis (Figure S1). So, for this analysis, we keep the following TCGA projects: KIRP, HNSC, COAD, LIHC, LUSC, LUAD, and BRCA. The age distribution of the two groups is shown in figure S2.

The differential expression and overlap analysis were done as before. To compare the expression (Fold Change) between the two groups, genes that lie in the overlap between the cancer-DEG and one of the four categories in each group were compared by Mann-Whitney U test, and the difference was considered significant when FDR <0.05.

## Results

### Loss of cellular identity in ageing identified in 40% of tissues analyzed

First, we identified cancer-DEG (tables S4-19) and ageing-DEG (tables S20-45) in a tissue-specific manner for 26 tissues and 16 cancer types. Moreover, we classified all the genes according to their tissue specificity based on tau’s categories (table S46). The number of DEGs and genes classified as pan-tissue or tissue-specific categories are shown in Table 1 for the cancers and tissues analyzed.

We then overlapped the ageing-DEGs with the Tissue-specific category (Figure 1A). We do not observe a clear pattern of loss (or gain) of specificity in a global manner. However, it is important to note that of the tissues analyzed, around 40% show enrichment of downregulated Tissue-specific genes (Figure 1A), with some exceptions. Adipose tissue, brain, colon, esophagus, lung, muscle, prostate, skin, small intestine, and testis present downregulation of Tissue-specific genes, without presenting significant results in the opposite direction. Validating these results, we observed similar patterns in genes with High specificity (Figure 2A). No significant results were identified for Tissue-unexpressed (Figure 2B).

**Figure 1:**
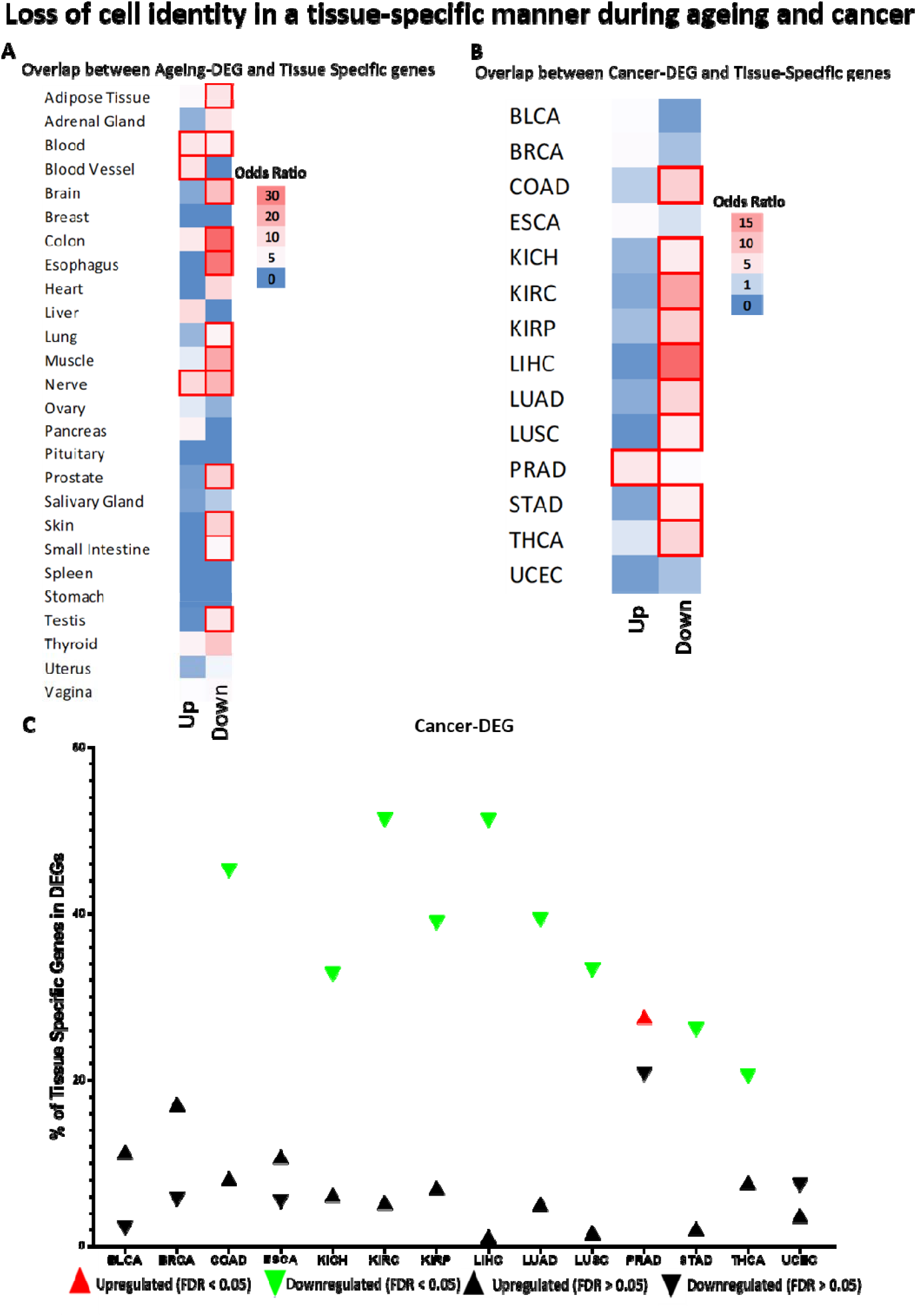
Overlap between DEGs and Tissue-Specific genes. A-B Heatmap of odds ratio on the chance of the overlap. Red borders represent significant results (FDR < 0.05). A Ageing-DEGs, B Cancer-DEGs. “Up” and “Down” represent whether genes are upregulated or downregulated. C Percentages of specificity-genes that are differentially expressed in cancer. Each dot represents the percentage of genes and the direction of expression. Coloured dots are the statistically significant results.

**Figure 2:**
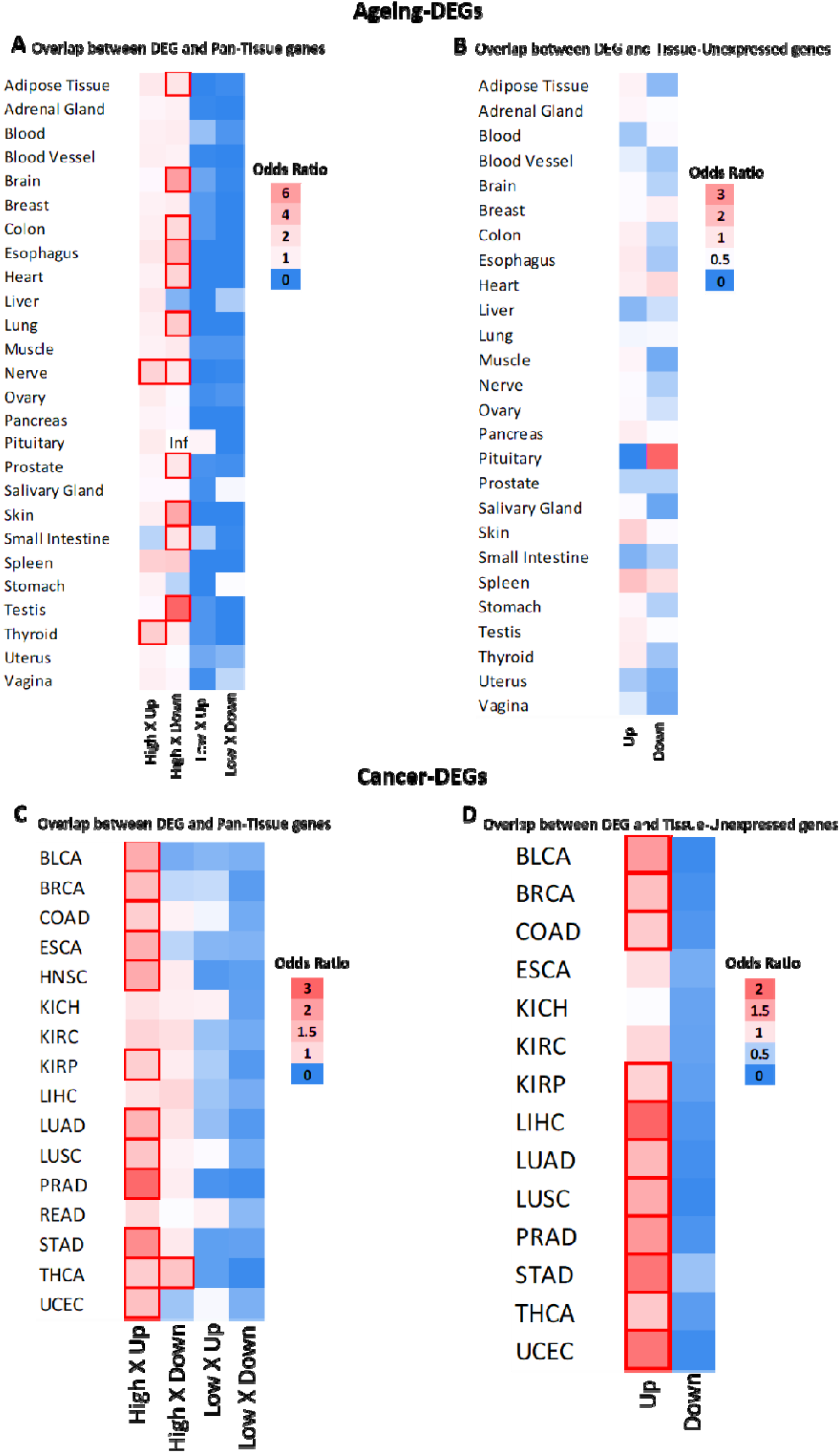
Overlap between DEGs and specificity categories. Heatmap of odds ratio on the chance of the overlap. Red borders represent significant results (FDR < 0.05). A-B Ageing-DEGs; C-D Cancer-DEGs. Inf = Odds ratio tends to infinity due to the low number of downregulated DEGs, but the result is not significant. “Up” and “Down” represent whether genes are upregulated or downregulated. “High” an “Low” represents Pan-Tissue group.

This body of results suggests that, although we have a tendency of loss of tissue identity for some tissues with ageing, it is not a global phenomenon, presenting some exceptions.

### A robust pattern of cellular identity loss is observed in most cancers studied

Repeating the same approach as before, we overlay the cancer-DEGs with the four specificity categories and observe a clear pattern in most of the cancers analyzed. First, we observed an enrichment of downregulated DEGs in Tissue-specific genes (Figure 1B). Validating these results, we can see a significant number of upregulated DEGs in High specificity and Tissue-unexpressed (Figure 2C-D, respectively).

Again, it is essential to highlight that the results of Tissue-specific genes (Figure 1B) are the most relevant, as opposed to the overexpression of genes from other organs and typically inactive genes (Figure 2C-D), which is expected considering the nature of cancer. We show in Figure 1C the percentage of Tissue-Specific genes that are cancer-DEGs, highlighting the results that were statistically significant from the previous analysis. We can clearly see a substantial proportion of Tissue-Specific genes downregulated in cancer, ranging from 20.5% in THCA to 51.3% in KIRC, considering statistically significant results.

This makes biological sense and is in line with the literature. Considering the pan-tissue group, our analyses showed cancers commonly overexpress genes associated with specific functions of other organs and tissues (High specificity genes). In a tissue-specific manner, they activate genes usually unexpressed and downregulate genes typically highly expressed in the original tissue. THCA (Figure 2C) and PRAD (Figure 1B and Figure 1C) are the only significant exceptions to this pattern.

Furthermore, as we found a trend of overexpression of High Specificity genes, we tried to answer which healthy tissues these genes are typically expressed in, repeating the same approach but overlapping only the upregulated cancer-DEGs with the Tissue-specific genes from all GTEx tissues. As shown in Figure S3, we found no clear pattern, which is aligned with the notion of generalized genetic instability in cancer cells.

To further validate our results, we repeated the analyses using an alternative gene classification (Figure S4). The biggest difference between the two analyses is in the results of the pan-tissue categories (Figures S4A and D). This is probably because the alternative categories are much less stringent, which results in many more genes and consequently more significant results. However, results from Tissue-specific group are quite similar (Figures S4B, C, E and F), indicating that the observed pattern is biologically relevant.

### Loss of tissue-identity is associated with cancer prognosis

After identifying the pattern of loss of tissue specificity in cancer, we sought to understand whether this impacts patient survival. To do this, we used the genes in the overlap between cancer-DEG and one of the four categories of tau specificity and built an expression signature. Using the median of expression as a cutoff, we constructed overall survival and disease-free survival curves comparing the high and low expression signature groups (Figure 3). Looking at the heatmaps (Figure 3A and B), we can see a trend in the Tissue-specific group. We can observe that most cancers show a positive hazard ratio pattern (i.e., high expression group associated with the worst survival) in the Tissue-unexpressed genes, and oppositely, a negative hazard ratio pattern (i.e., low expression group associated with the worst survival) in Tissue-specific genes.

**Figure 3:**
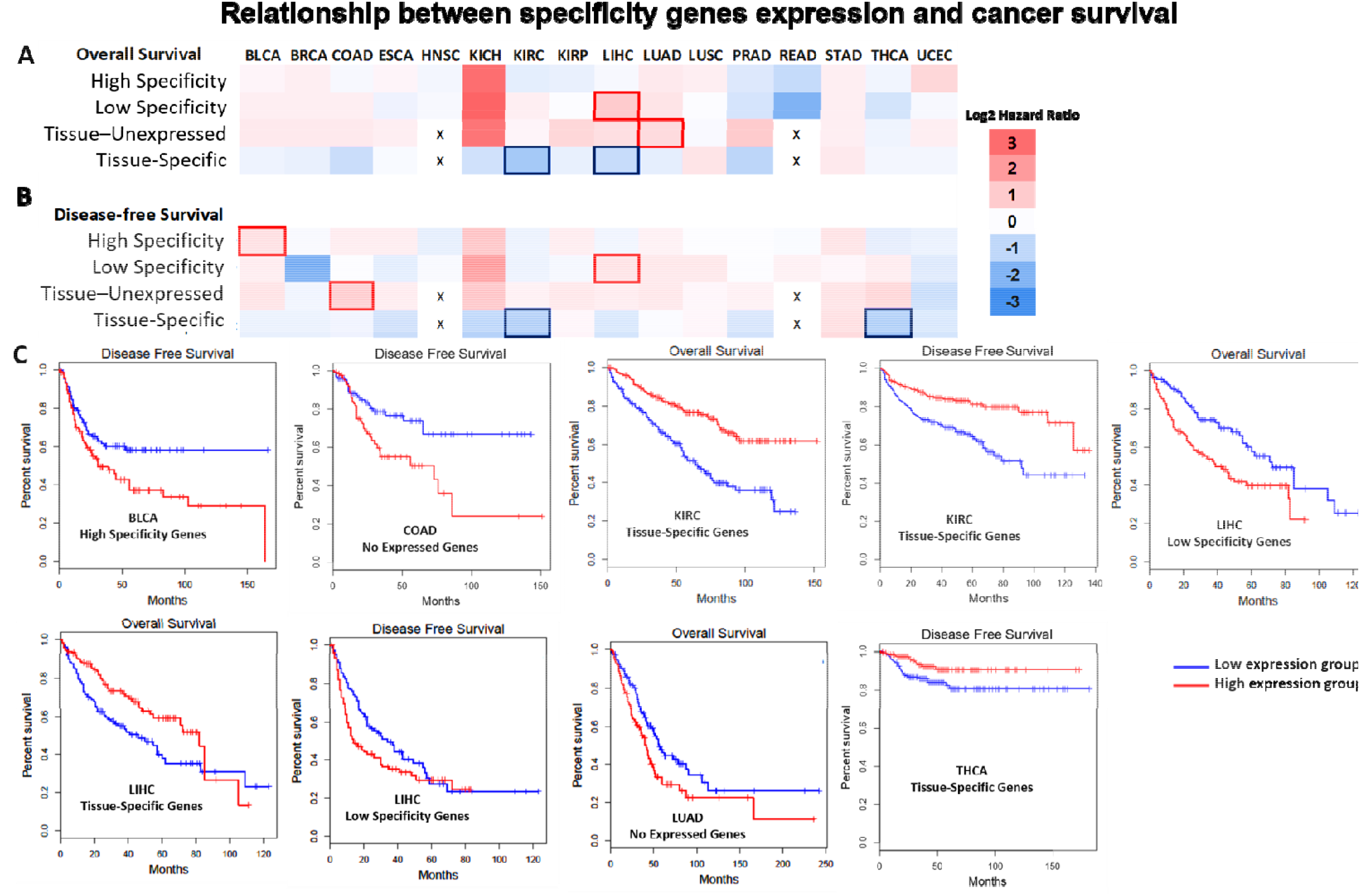
Relationship between specificity genes and cancer survival. A and B Heat map of hazard ratio of overall and disease-free survival, respectively, statistically significant results (Mantel-Cox test, FDR < 0.1) are highlighted with blue or red borders, according to the direction of the expression signature and the worst survival. The x represents where analysis cannot be done in a tissue-specific manner. C) Kaplan-Meier curves of the significant results of the previous heatmaps.

When analyzing the curves of the significant results (Figure 3C), we observe that almost all observations align with the pattern observed previously: The upregulation of High specificity or Tissue-unexpressed is associated with worsened survival, and the downregulation of Tissue-specific genes is associated with worse prognosis. A partial exception is observed in LIHC, where the overexpression of Low specificity genes is related to a worsening in overall and disease-free survival. Considering these results, we suggest a trend for the loss of tissue identity to increase cancer aggressiveness.

### Age does not influence the loss of tissue specificity in cancer

Previous studies demonstrated that there are important molecular differences when considering the age of cancer patients (15, 31-33). Then, considering that perhaps the loss of tissue identity occurs in ageing, we tested the hypothesis that the patient’s age influences the pattern of expression of specificity genes. For this, we separated the TCGA cancers into two groups in relation to age, as described in the methods, and generated two lists of DEGs for each tumor (old cancer-DEG young cancer-DEG). Old cancer-DEGs and young cancer-DEGs are in Tables S47-60.

We then performed the overlap between the DEGs of the two groups (Table 2) and observed that most differentially expressed genes are shared independent of patient age, but a few hundred genes are unique to the old or young group. Besides that, 5 genes showed opposite expression patterns: COX4I2 in HNSC; NR4A2 and NR4A1 in COAD; and CYP26A1 and FDCSP in BRCA. It would be interesting to explore whether these genes are important in differentiating cancers from old and young patients, but this analysis is beyond the scope of this study.

**Table 2.** Overlap between old and young cancer-DEGs. *”Up” and “Down” represent whether genes are upregulated or downregulated

|  | Up-Old and Young<br>Shared | Down-Old and Young<br>Shared | Up-Old<br>Exclusive | Down-Old<br>Exclusive | Up-Young<br>Exclusive | Down-Young<br>Exclusive |
| --- | --- | --- | --- | --- | --- | --- |
| **KIRP** | 1176 | 777 | 210 | 220 | 784 | 434 |
| **HNSC** | 866 | 670 | 310 | 324 | 323 | 604 |
| **COAD** | 1406 | 1276 | 393 | 391 | 183 | 138 |
| **LIHC** | 1108 | 553 | 226 | 156 | 523 | 335 |
| **LUSC** | 2301 | 1837 | 247 | 338 | 424 | 236 |
| **LUAD** | 1616 | 1132 | 170 | 354 | 438 | 314 |
| **BRCA** | 1484 | 1110 | 568 | 514 | 441 | 158 |
| **Total** | 9957 | 7355 | 2124 | 2297 | 3116 | 2219 |

Focusing on the main objective of this study, we repeated the overlap analysis but now considering the age groups (Figure 4A-C). We can observe that the pattern is, in general, the same as previously observed, and age does not change it. Next, we directly compared the expression (fold change) of the genes of interest in relation to the four specificity categories (Figure 4D-G). Although we have a few significant differences, they are slight and in the same direction, making it clear that age is not influencing the loss of cancer tissue identity, reinforcing this phenomenon may be essential for carcinogenesis.

**Figure 4:**
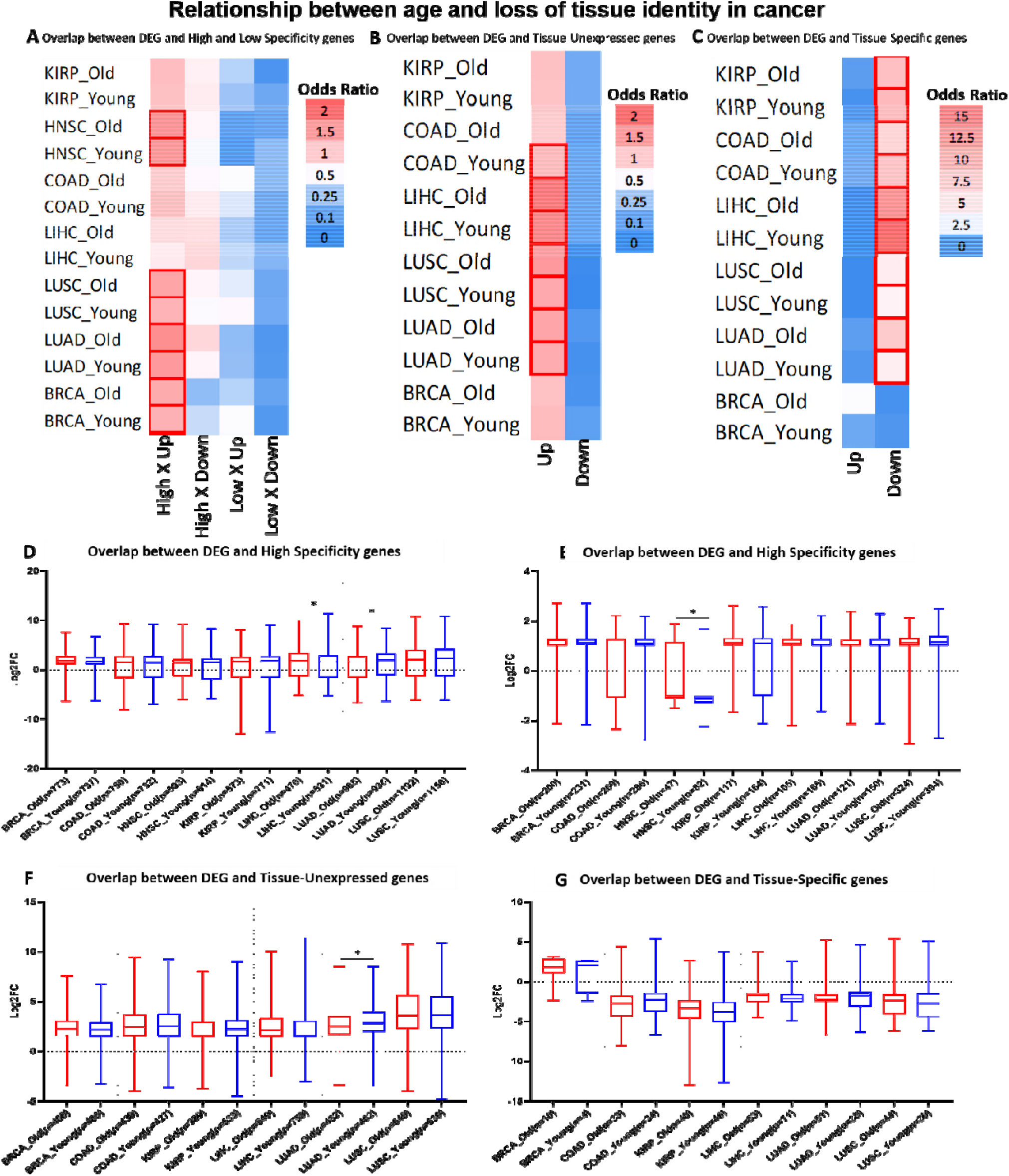
Overlap between cancer-DEGs and specificity categories considering patient age groups. A-C, heatmap of odds ratio on the chance of the overlap. Red borders represent significant results (FDR < 0.05). D-G, plots comparing the expression level (Fold Change) of the overlap genes between DEGs and specificity categories in the two age groups (red = old and blue = young) significant results are represented with * (FDR < 0.05). Error bars represent the maximum and minimum values. Number of genes in each group is in parentheses on the x-axis. “Up” and “Down” represent whether genes are upregulated or downregulated

## Discussion

Cell identity and plasticity is an essential topic in oncology and, more recently, is gaining importance in gerontology (8, 34, 35). In this work, we analyze how changes in gene expression are related to tissue specificity during cancer and ageing, using data from thousands of human samples.

First, we sought to test the hypothesis that tissues lose their identity “naturally” in ageing. Although we see a trend of downregulation of Tissue-Specific genes (which could reinforce the hypothesis), it occurred in less than half of the tissues, suggesting that, at least, this phenomenon is not valid for the whole organism or is too subtle to be detected amid all the transcriptional noise in ageing, especially from bulk RNA-seq data (36). Izgi et al., observed a loss of cellular identity in brain, lung, liver, and muscle in ageing mice, our results suggest similar findings only in brain, lung and partially in muscle (4). In the same paper, the authors also analyzed GTEx data, and similar to our study, they did not find a clear pattern of inter-tissue convergence during ageing in humans.

In cancer, on the other hand, we simultaneously observe an upregulation of High specificity genes, downregulation of Tissue-specific genes, and activation of Tissue-unexpressed genes. This suggests that during tumorigenesis, cancer cells gain functions of other organs/tissues (or at least there are more upregulated genes because of the noise from generalized genome instability) while suppressing the functions of their original tissue. Besides making sense biologically, this result aligns with the literature since dedifferentiation is a known feature of cancer (37, 38). This process has been described in some individual cancers such as colon, melanoma, and pancreas, but as far as we know, we were the first to demonstrate this in a pan-cancer analysis and in a tissue-specific manner (39-41).

Exploring cancer as a tissue-specific disease is an approach that is gaining prominence in oncology, with several studies trying to understand in depth the genetics that regulate this process (10, 42, 43). Schaefera et al. explore why some genetic alterations are only relevant in specific types of cancer, concluding that the tissue microenvironment is a determining factor in this process (44). Two other studies have demonstrated that expression signatures can help classify the tissue for cancers of unknown primary origin, which presents a possible application of using the transcriptome signatures with tissue specificity in oncology (45, 46). Our work, besides adding novel knowledge to this field, corroborates studies such as that from Hu et al., which showed that in cancer, there is a decrease in the expression of some tissue-specific genes, and Pei et al., which showed that it is common for cancers to acquire specific expression profiles from other organs (47, 48).

When we directly compare cancer results with ageing, we have an interesting finding: in cancer, we have an upregulation of High specificity genes, and in ageing a trend to downregulation. This kind of opposite pattern is expected and has already been described by our group (3). But when analyzing the Tissue-Specific genes, most of the significant results are in the same direction, with the downregulation of these genes. This makes us wonder if preventing the loss of tissue specificity might be a promising strategy against cancer and ageing at the same time. But this result needs to be looked at carefully since less than half of the normal tissues show this pattern, and there are a few exceptions.

The pattern found also seems to influence the aggressiveness of cancer, impacting on patient survival. A phenomenon linked to the loss of tissue specificity is the acquisition of stemness and dedifferentiation, which is also related to the aggressiveness of cancers (38). Furthermore, we still need better biomarkers in oncology, and although our data needs to be refined for possible application, tissue identity loss has the potential to improve the prognostic classification of cancer patients (49-51).

Finally, we tested the hypothesis that the age of patients influences expression patterns of specificity genes. We found no relevant difference between the young and old groups, indicating that age does not affect the process of tissue identity loss in cancer. This reinforces the newly proposed idea that the phenomenon of acquiring cellular plasticity (which includes loss of identity) is a hallmark of cancer (7).

In summary, we show evidence of an age-dependent loss of tissue specificity in some tissues; however, this process is not global, nor are our results robust enough to confirm the hypothesis of loss of cell identity in ageing. On the other hand, in cancer, we have a pattern of clear downregulation of Tissue-specific genes and activation of genes not expressed in the original tissue, including genes highly expressed in other tissues. Our results also suggest that this pattern influences cancer aggressiveness and is not influenced by the patient’s age, corroborating that it is a crucial step for carcinogenesis.

## Acknowledgements

We are grateful to current and past members of the Genomics of Ageing and Rejuvenation Lab for valuable discussions.

## Supplementary Materials

**Figure S1:**
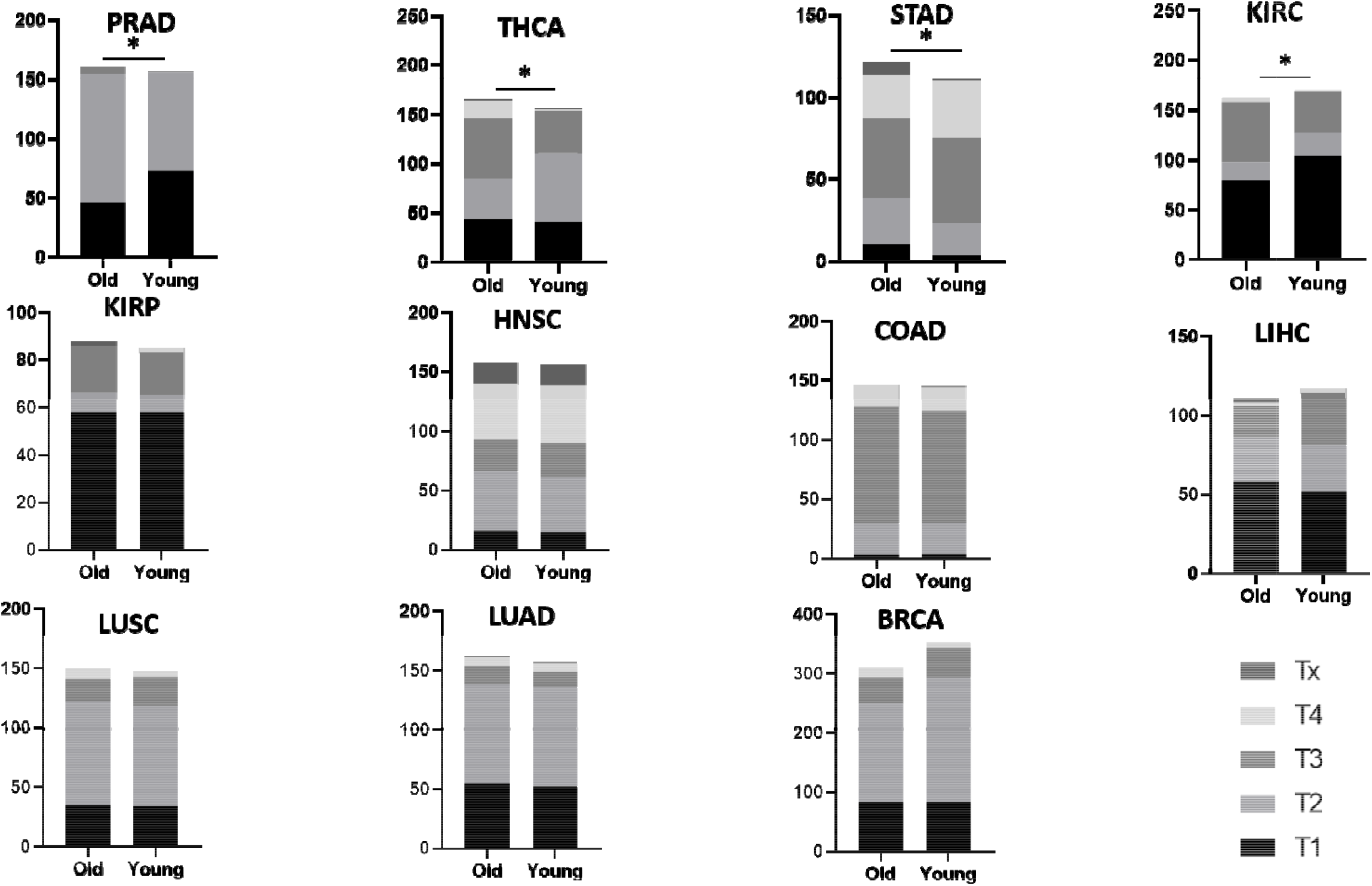
Comparison between the T pathological stage proportions of the old and young groups. The y-axis represents the number of patients. * p-value<0.05

**Figure S2:**
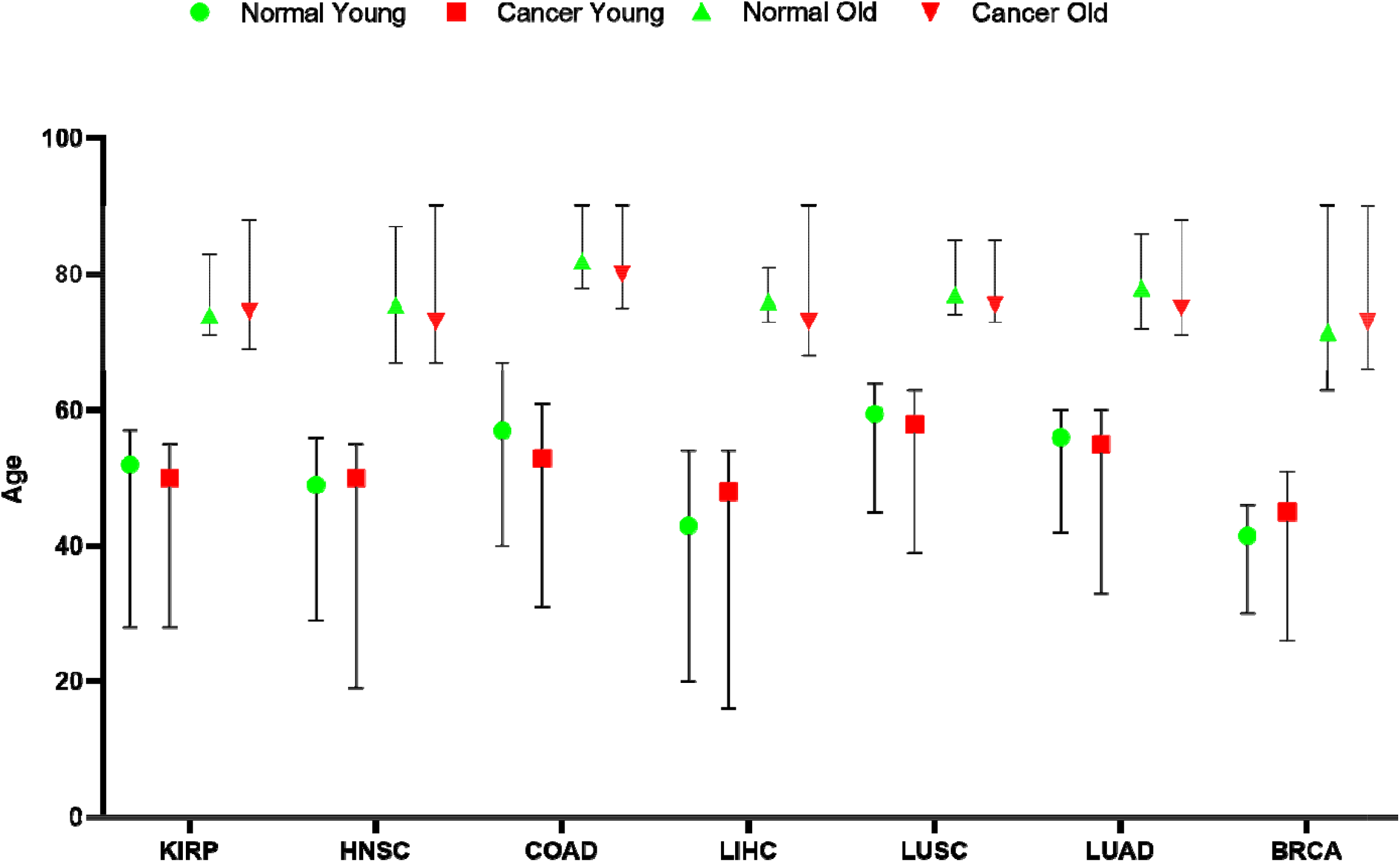
Age distribution of the seven cancers used in the age group analysis. Error bars represent the maximum and minimum values.

**Figure S3:**
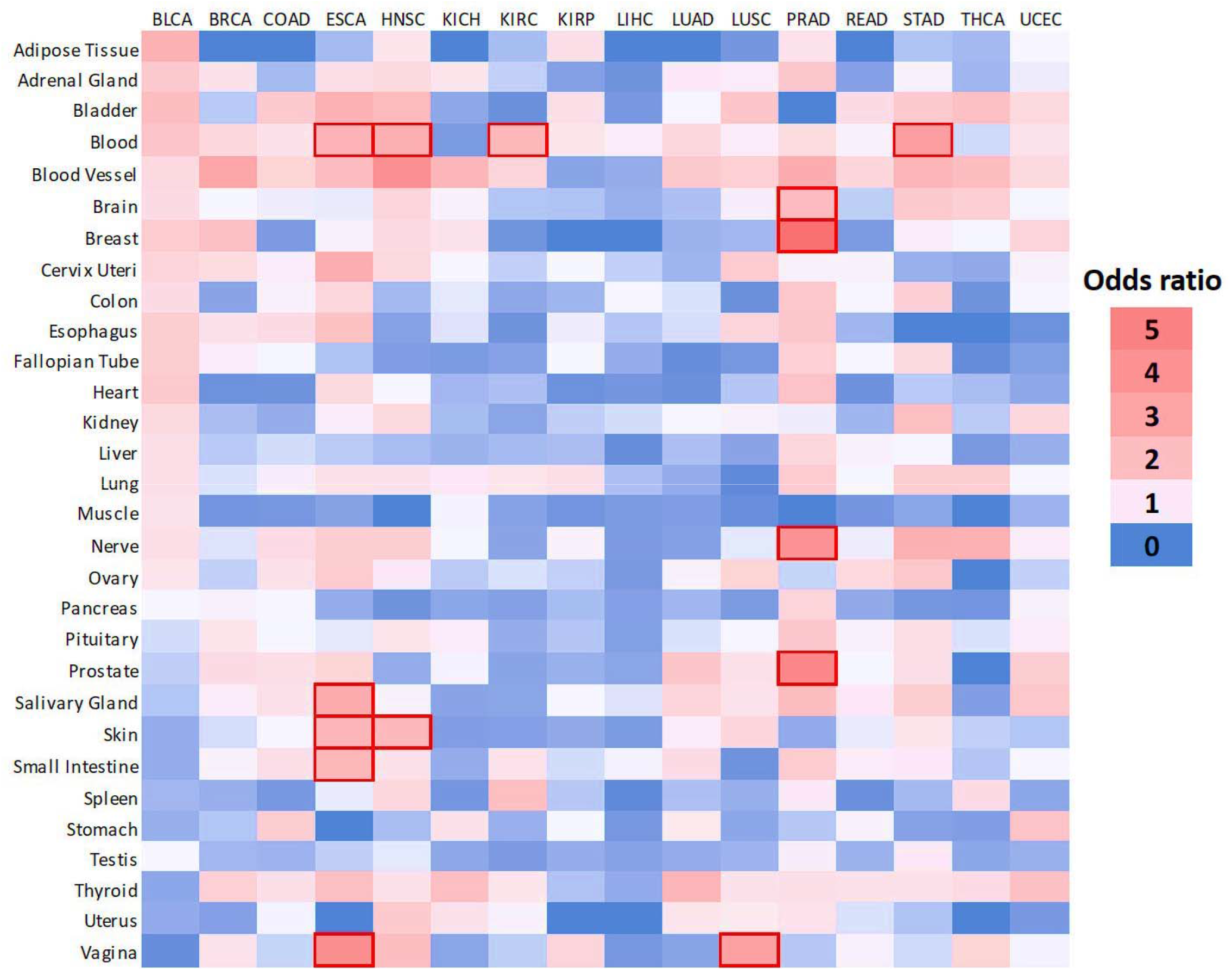
Overlap between upregulated cancer-DEGs and Tissue-Specific genes. Heatmap of odds ratio on the chance of the overlap. In the columns we have the TCGA cancers and in the rows the GTEx tissues. Red borders represent significant results (FDR < 0.05).

**Figure S4:**
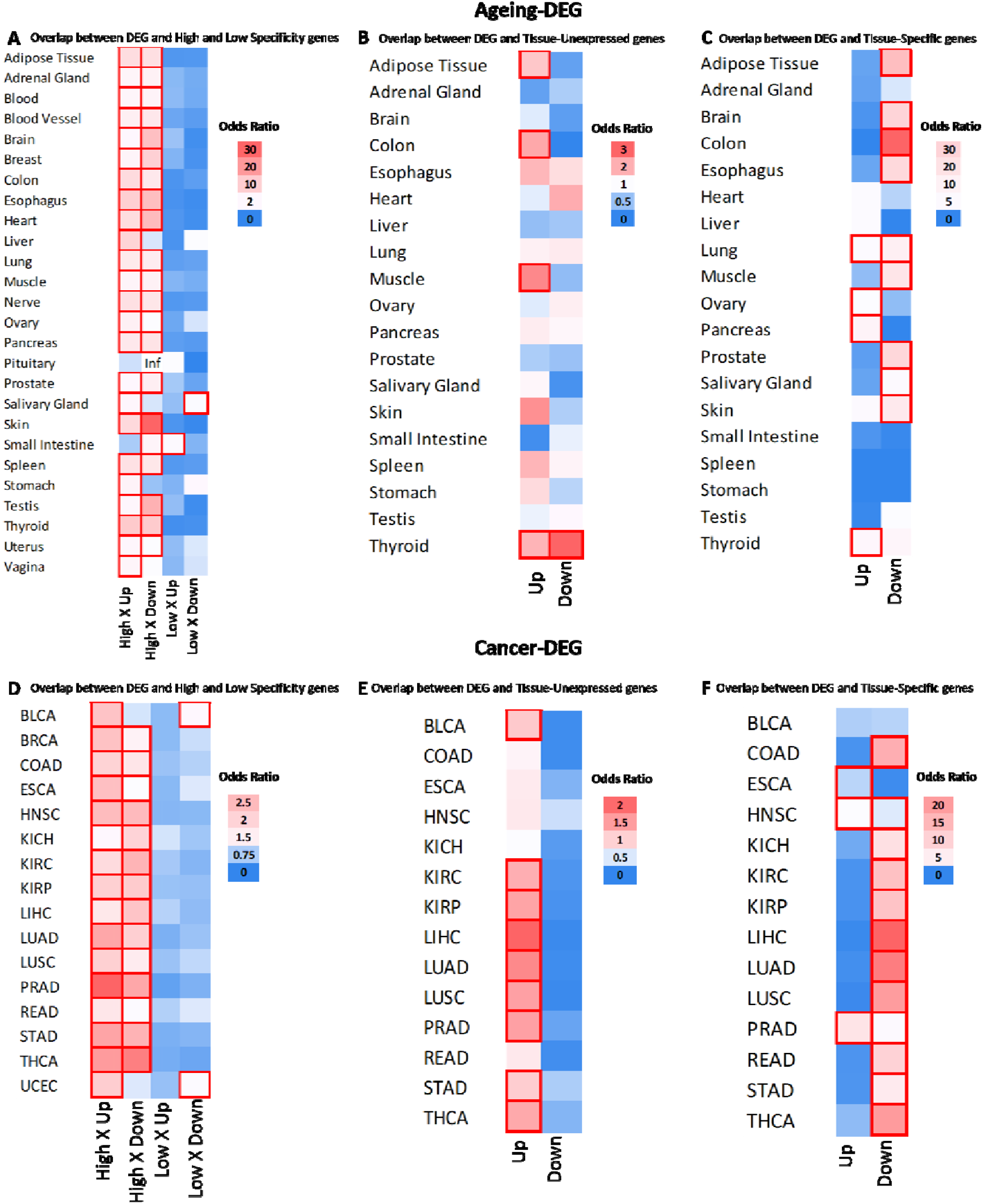
Overlap between DEGs and alternative specificity categories. Heatmap of odds ratio on the chance of the overlap. Red borders represent significant results (FDR < 0.05). A-C Ageing-DEGs, D-E Cancer-DEGs. Inf = Odds ratio tends to infinity due to the low number of downregulated DEGs, but the result is not significant. “Up” and “Down” represent whether genes are upregulated or downregulated.

